# Combined Metabolic Activators decrease liver steatosis by activating mitochondrial metabolism in a Golden Syrian hamster study

**DOI:** 10.1101/2021.02.19.431968

**Authors:** Hong Yang, Jordi Mayneris-Perxachs, Noemí Boqué, Josep M del Bas, Lluís Arola, Meng Yuan, Hasan Turkez, Mathias Uhlén, Jan Borén, Zhang Cheng, Adil Mardinoglu, Antoni Caimari

**Affiliations:** Science for Life Laboratory, KTH - Royal Institute of Technology, Stockholm, Sweden; Department of Diabetes, Endocrinology and Nutrition, Hospital Universitari de Girona Doctor Jospe Trueta and Girona Biomedical Research Centre (IDIBGI), Girona, Spain; CIBER Pathophysiology of Obesity and Nutrition (CIBEROBN), Instituto de Salud Carlos III, Madrid, Spain; Eurecat, Centre Tecnològic de Catalunya, Technological Unit of Nutrition and Health, Reus, Spain; Universitat Rovira i Virgili, Nutrigenomics Research Group; Department of Biochemistry and Biotechnology, Campus Sescelades, Tarragona, Spain; Department of Medical Biology, Faculty of Medicine, Atatürk University, Erzurum, Turkey; Department of Molecular and Clinical Medicine, University of Gothenburg and Sahlgrenska University Hospital, Gothenburg, Sweden; School of Pharmaceutical Sciences, Zhengzhou University, Zhengzhou, PR China; Centre for Host-Microbiome Interactions, Faculty of Dentistry, Oral & Craniofacial Sciences, King’s College London, London, United Kingdom

## Abstract

The prevalence of non-alcohol fatty liver disease (NAFLD), defined as the liver’s excessive fat accumulation, continues to increase dramatically. We have recently revealed the molecular mechanism underlying NAFLD using in-depth multi-omics profiling and identified that combined metabolic activators (CMA) could be administered to decrease the amount of hepatic steatosis (HS) in mouse model and NAFLD patients based on systems analysis. Here, we investigated the effects of a CMA including L-carnitine, N-acetyl-l-cysteine, nicotinamide riboside and betaine on a Golden Syrian hamster NAFLD model fed with HFD, and found that HS was decreased with the administration of CMA. To explore the mechanisms involved in the clearance HS, we generated liver transcriptomics data before and after CMA administration, and integrated these data using liver-specific genome-scale metabolic model of liver tissue. We systemically determined the molecular changes after the supplementation of CMAs and found that it activates mitochondria in the liver tissue by modulating the global fatty acid, amino acids, antioxidant and folate metabolism.

## Introduction

Hepatic steatosis (HS) is defined as the excessive accumulation of fat in the liver (>5.5% tissue weight), and it is the most common liver disease worldwide ^1^. Non-alcoholic fatty liver disease (NAFLD), a widespread metabolic disorder, refers to a group of conditions including HS and various degrees of liver inflammation such as non-alcoholic steatohepatitis (NASH). NAFLD can progress to cirrhosis, and ultimately hepatocellular carcinoma (HCC) which are much more severe liver diseases ^1, 2, 3^. It has been reported that the global prevalence of NAFLD is approximately at ∼25% ^1, 4, 5^. Although research in drug development for NAFLD is intense and advancing rapidly, there are still significant unmet challenges with no effective drug approved for this condition ^6^.

The insufficient capacity to remove incomplete products of fatty acid oxidation is one of NAFLD’s main hallmark ^7^. Oxidative stress has a significant role in NAFLD’s pathogenesis and is characterized by impaired function of electron transport chains, impaired oxidation and increased production of reactive oxygen species (ROS) ^8, 9, 10^. Glutathione (GSH) is the most abundant endogenous antioxidant in response to oxidative stress and increased ROS ^11, 12, 13^. Our previous studies, combing personalised genome-scale metabolic models (GEMs) and in-depth multi-omics profiling based on clinical data indicated an augmented requirement for nicotinamide adenine dinucleotide (NAD^+^) and GSH in NAFLD patients. A three-step strategy including (i) activating mitochondrial fatty acid uptake, (ii) increasing fatty acid oxidation, (iii) increasing the availability of GSH can be employed to decrease of the HS in NAFLD patients^14, 15, 16, 17, 18^.

To validated our hypothesis, we performed a study in which supplementation of a CMA including serine (GSH precursor), nicotinamide riboside (NR, NAD^+^ precursor) and N-acetyl-l-cysteine (NAC, GSH precursor) was given to mice fed a high-fat diet (HFD) for 14 days. The results showed that supplementation of CMA significantly decreased the amount of liver fat by promoting the fat oxidation in mitochondria in the liver ^15, 19^. We further studied the effect of the CMA including serine, NR, NAC and L-carnitine tartrate (LCT, salt form of l-carnitine that promote the update of fat to mitochondria) in a one-day proof-of-concept human supplementation study (*NCT03838822*). We observed that the supplementation resulted in increased of fatty acid oxidation and *de novo* GSH synthesis ^20^. Recently, we also tested the effect of the long-term effect of the CMA including serine, NR, NAC and LCT in 10 weeks randomised, placebo-controlled study and found that administration of CMA decreases the amount of HS in liver together with other clinical parameters including AST, ALT, uric acid and creatinine ^21^. However, the underlying molecular mechanism associated with decreased HS has not been revealed in animal or human studies.

In this study, we investigated the effect of the supplementation of CMA, including NR, L-carnitine, NAC, and betaine (as a GSH precursor since that can be converted to glycine and serine), in Golden Syrian hamsters with NAFLD induced by HFD. We used Golden Syrian hamsters since it is a better model for experimental studies on lipoprotein metabolism than other smaller-sized animals, e.g., rat and mouse ^22, 23, 24^. Furthermore, we previously used this model to evaluate successfully the health effects of different bioactive compounds such as phytosterols ^25^ and extracts of hazelnut skin ^26^ and grape seed procyanidins ^25, 27^ against fatty liver, dyslipidaemia and adiposity. In the present study, we found that administration of CMA ameliorated HS. To explore the underlying molecular mechanisms involved in decreased HS, we generated liver transcriptomics data before and after supplementation of CMA and integrated these data using liver GEM. Finally, we systematically identified the phenotypical differences and presented the potential effects of CMA supplementation on the global metabolism based on systems analysis.

## Results

### CMA attenuate HFD-induced HS in a Golden Syrian hamster NAFLD model

We fed Golden Syrian hamsters with either normal fat diet (NFD) or HFD for 8 weeks. Afterwards, we provided either a placebo or a CMA including NR, L-carnitine and NAC by *la gavage* and betaine in the drinking water to hamsters at intended human clinical doses (Dose I; HFD+MI_D1 group) and 2-fold (Dose II; HFD+MI_D2 group) dose levels for 4 weeks (Figure 1A). We assessed the changes in body weight, biometric and serum variables of each hamster group at week 8 and 12 and examined the phenotypic differences in all groups. We observed that the hamsters fed with HFD had significantly more severe grade of HS compared to those from the NFD-fed group (p < 0.05), and this HFD-induced HS was significantly attenuated in CMA-treated hamsters with Dose I and II (Figure 1B). Additionally, the histological analyses showed a significant increase of lipid deposition in the HFD group’s liver compared to those of the control group (Figure 1C&D). Specifically, the HFD groups developed microvesicular steatosis without apparent inflammation, sinusoidal dilatation or fibrosis (Figure 1D), and CMA supplementation with Dose I and II reduced the HFD-induced lipid deposition in the liver (Figure E&F). The HFD-fed hamsters, compared to the control group, showed significantly increased liver weights (p = 0.001) and relative mesenteric adipose tissue depots (MWAT%, mean increase 37.5%, p = 0.009) (Table 1). CMA treatment tended to induce a reduction in liver weight, mainly in hamsters supplemented with Dose II (8.8% and 10.1% reduction with Dose I and II, respectively) compared to the HFD group. (Figure 1G). Body composition analysis revealed that HFD feeding led to a weight gain during the last 4 weeks of the study (mean increase 4.5%, p = 0.03) (Figure 1H, Table 1). Hamsters treated by CMA with Dose II showed significantly (p = 0.005) reduced body weight (5.1% mean reduction) (Figure 1H, Table 1) and significantly (p = 0.05) increased lean/fat mass ratio (10.9% mean increase) (Figure 1I, Table 1) between 8 weeks and 12 weeks. Furthermore, at the end of the study, the supplementation with Dose II significantly decreased body weight (p = 0.021) and body weight gain (p = 0.0004) compared to the HFD-fed hamsters that received the placebo (Figure 1H, Table 1). This body weight loss could be attributed, at least in part, to the decreased cumulative caloric intake (p = 0.009) displayed by the HFD+MI_D2 group during the 4-week CMA supplementation period when compared to the HFD group. A similar trend in body weight (p = 0.137) and body weight gain (p = 0.066) was observed in hamsters treated with Dose I in comparison with HFD group, although the differences were not significant (Figure 1H, Table 1). We also measured the NAD^+^ levels in the liver and found that its levels significantly increased with CMA Dose I and II supplementation compared to their HFD counterparts, as predicted in our earlier studies (Figure 1J).

**Figure 1.**
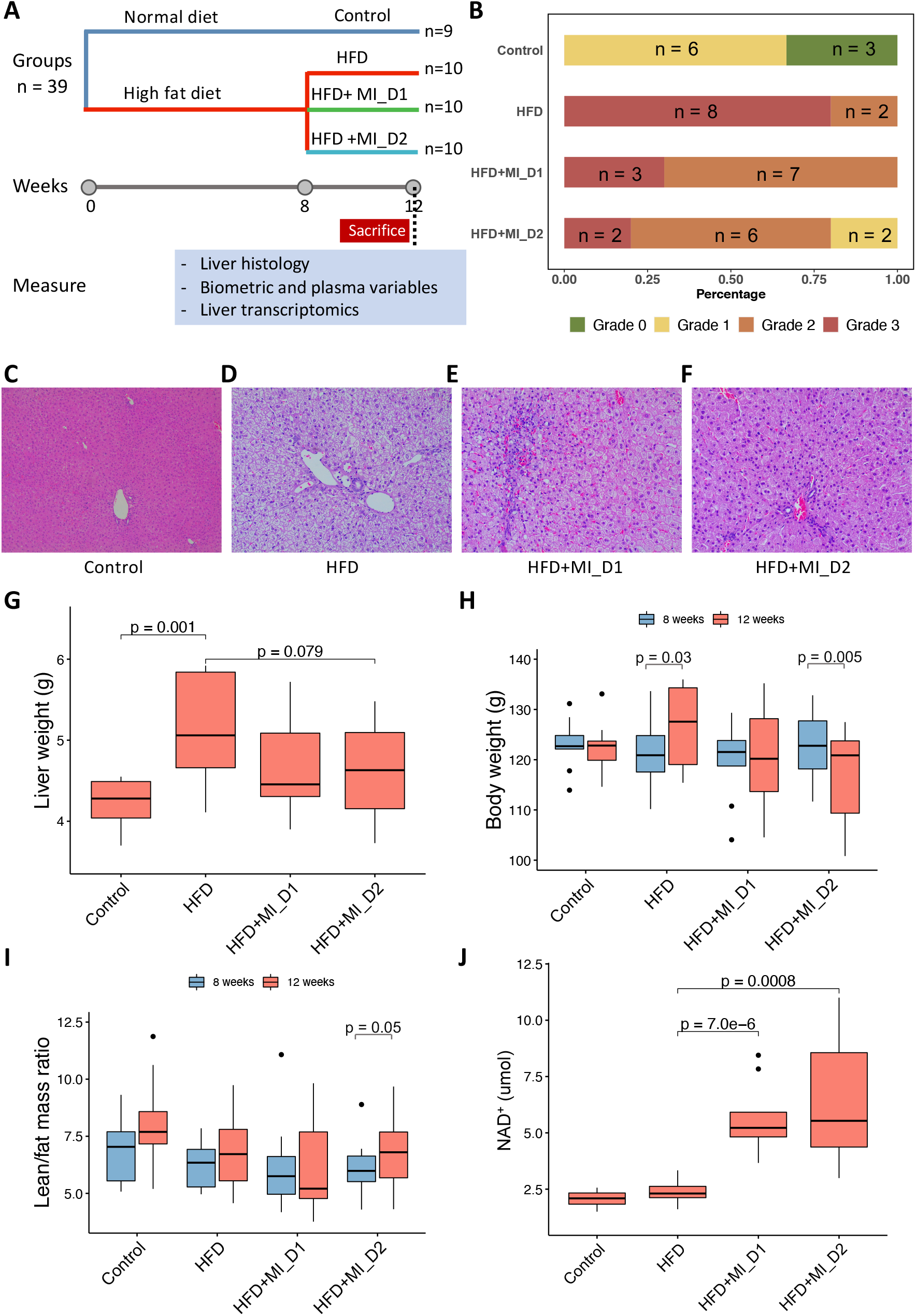
Effect of HFD and CMA treatment on liver histology, liver weights and body weight of Golden Syrian hamsters. (a) Study design. (b) Steatosis score of histological changes in liver. (c-f) Histological analysis of steatosis in liver sections stained with H&E of liver. (g) Liver weight. (h) Body weight. (i) Lean / fat mass ratio. (j) The hepatic expression of NAD^+^. Control, normal diet; HFD, high-fat diet; HFD+MI_D1, high-fat diet supplemented with CMA at Dose I; HFD+MI_D2, high-fat diet supplemented with CMA at Dose II.

**Table 1:** Effects of HFD and multi-ingredients treatment on biometric and serum variables.

|  | Control<br>(n = 9) | HFD<br>(n = 10) | HFD + MI_D1<br>(n = 10) | HFD + MI_D2<br>(n = 10) |
| --- | --- | --- | --- | --- |
| **Biometric variables** |  |  |  |  |
| Initial body weight (g) | 123.05 ± 5.14 | 121.07 ± 7.54 | 119.85 ± 7.51 | 122.55 ± 6.90 |
| Final body weight (g) | 122.20 ± 5.62 | 126.57 ± 8.20<sup>d</sup> | 120.08 ± 10.32 | 116.28 ± 9.86<sup>cf</sup> |
| Liver weight (g) | 4.23 ± 0.30 | 5.14 ± 0.65<sup>a</sup> | 4.69 ± 0.60 | 4.62 ± 0.61 |
| Liver weight (%) | 3.49 ± 0.25 | 4.04 ± 0.29<sup>a</sup> | 3.92 ± 0.43 | 3.98 ± 0.29 |
| Initial fat mass (%) | 11.94 ± 3.20 | 13.87 ± 1.95 | 14.40 ± 3.12 | 14.04 ± 2.28 |
| Final fat mass (%) | 11.28 ± 2.60 | 13.14 ± 2.78 | 14.84 ± 4.19 | 13.10 ± 2.87 |
| Initial lean mass (%) | 84.89 ± 3.26 | 84.27 ± 1.94 | 84.06 ± 2.99 | 84.02 ± 2.20 |
| Final lean mass (%) | 85.42 ± 2.40 | 85.74 ± 3.03 | 83.95 ± 4.04 | 85.22 ± 3.09<sup>f</sup> |
| Initial lean/fat mass ratio | 7.83 ± 3.22 | 6.20 ± 1.01 | 6.21 ± 1.97 | 6.16 ± 1.22 |
| Final lean/fat mass ratio | 8.00 ± 2.13 | 6.84 ± 1.67 | 6.20 ± 2.19 | 6.83 ± 1.70 |
| Cumulative food intake (kcal) | 126.77 ± 14.43 | 111.08 ± 13.22<sup>ad</sup> | 104.22 ± 15.94<sup>e</sup> | 95.23 ± 10.69<sup>cf</sup> |
| **Plasma variables** |  |  |  |  |
| Glucose (mM) | 7.29 ± 1.50 | 9.43 ± 2.01<sup>a</sup> | 8.93 ± 1.42 | 9.27 ± 2.01 |
| Insulin (mIU/L) | 16.69 ± 2.92 | 16.26 ± 3.61 | 16.17 ± 1.34 | 16.78 ± 3.43 |
| CHOL (mM) | 2.87 ± 0.21 | 4.03 ± 0.65<sup>a</sup> | 3.77 ± 0.60 | 4.44 ± 0.85 |
| HDL-C (mM) | 2.57 ± 0.37 | 3.15 ± 0.50<sup>a</sup> | 2.88 ± 0.52 | 3.32 ± 0.60 |
| LDL-C (mM) | 0.90 ± 0.20 | 1.38 ± 0.33<sup>a</sup> | 1.22 ± 0.24 | 1.34 ± 0.28 |
| TG (mM) | 1.07 ± 0.30 | 1.15 ± 0.53 | 0.96 ± 0.31 | 0.82 ± 0.32 |
| RWAT (%) | 1.03 ± 0.23 | 0.87 ± 0.17 | 0.99 ± 0.24 | 0.90 ± 0.13 |
| MWAT (%) | 0.88 ± 0.24 | 1.21 ± 0.24<sup>a</sup> | 1.17 ± 0.29 | 1.14 ± 0.19 |
| MUS | 0.27 ± 0.03 | 0.25 ± 0.03 | 0.24 ± 0.03 | 0.24 ± 0.03 |

Data are showed as the mean ± SD. Different superscript letters indicate statistically significant differences ( $p < 0.05$ , Student' t-test). a, b and c indicate difference between HFD and control groups; between HFD+MI\_D1 and HFD group; between HFD + MI\_D2 and HFD groups, respectively. d, e and f show difference between 8 weeks (initial) and 12 weeks (final) in HFD, HFD+MI\_D2 and HFD+MI\_D2 group, respectively. RWAT, retroperitoneal white adipose tissue. MWAT, mesenteric white adipose tissue. MUS, gastrocnemius and soleus muscles.

We observed reduction in plasma triglyceride concentrations in CMA-treated hamsters (16.5% and 28.7% mean reduction with Dose I and Dose II, respectively) (Table 1) than those in the HFD group in parallel to the decrease in HS. Additionally, we found that HFD significantly increased the plasma level of glucose (p = 0.02), total cholesterol (CHOL; p = 0.0001), high-density lipoprotein cholesterol (HDL-C; p = 0.01), and low-density lipoprotein cholesterol (LDL-C; p = 0.001) during the 12 weeks feeding period (Table 1). We observed that CMA supplementation with different doses did not alter these parameters (Table 1).

### Transcriptomics alteration with HFD and CMA treatment

To reveal the underlying molecular changes associated with the decreased HS after CMA treatment, we performed global transcriptomics analysis using RNA sequencing (RNA-seq) on liver tissue of each group. Globally, PCA revealed patterns of gene expression changes with HFD and CMA treatments (Figure 2A). At a 10% false discovery rate (FDR), we found 3,317 differentially expressed genes (DEGs) between HFD and control group, 1,669 of which were significantly up-regulated and 1,648 of which were down-regulated (Figure 2B, Dataset S1). We performed KEGG pathway enrichment analysis on up-regulated and down-regulated DEGs separately in each group. Our results indicated that up-regulated DEGs were enriched in pathways including oxidative phosphorylation, ribosome, metabolic pathway, lysosome, cardiac muscle contraction, and spliceosome in HFD group vs control group (Figure 2C, Dataset S2). We found that fatty acid metabolism, catabolism of amino acids including branched-chain amino acids (BCAAs) and lysine, metabolism of amino acids (tryptophan, beta-alanine, glycine, serine, and threonine), as well as signalling pathways in the regulation of insulin resistance and fatty acid oxidation (*e*.*g*., AMPK-signalling pathway) were significantly enriched with the down-regulated DEGs in HFD group vs control group (Figure 2C, Dataset S2).

**Figure 2.**
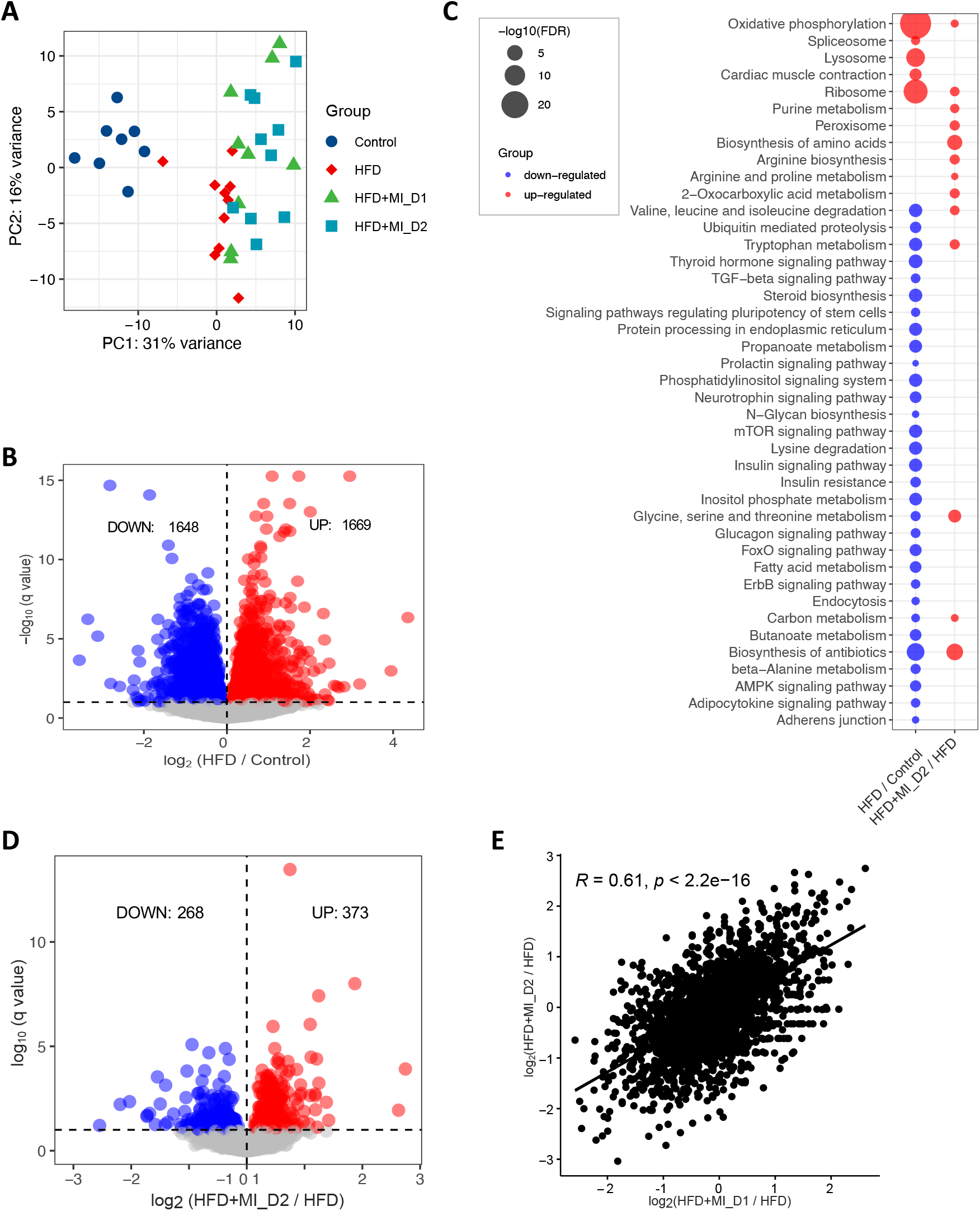
Effects of HFD and CMA on liver transcriptomics of Golden Syrian hamsters. (a) PCA plot of 37 hamsters. (b) Up- and down-regulated genes in HFD group as compared to control group. (c) KEGG pathway analysis shows pathways that were significantly altered between HFD and control groups, HFD+MI_D2 and HFD groups, respectively. Pathways that are down-regulated or up-regulated are shown in blue and red, respectively. The size of the bubble is proportional to -log_10_ of the FDR for each KEGG pathway term. (d) Up- and down-regulated genes in HFD+MI_D2 group as compared to HFD group. (e) The log 2-fold changes in differential expressed genes between two groups compared to HFD group is significantly positive correlated.

We also performed Gene Ontology (GO) enrichment analysis for the DEGs. Compared to the control group, we found that up-regulated genes in HFD group were most significantly enriched in nucleoside monophosphate metabolic process, ATP synthesis coupled electron transport, respiratory electron transport chain and purine nucleoside metabolic process. We also found that down-regulated genes were most significantly enriched in cellular protein modification process, protein modification process and cellular protein metabolic process in HFD vs control group (Dataset S2).

Next, we compared the gene expression changes between HFD and CMA-treated groups (fed with HFD + MI_D1 and HFD + MI_D2). In total, 80 (56 up-regulated and 24 down-regulated) and 641 DEGs (268 down-regulated and 373 up-regulated) (Figure 2D, Dataset S1) were identified in Dose I and II groups, respectively. We found that the log 2-fold changes in DEGs between two groups compared to HFD group are significantly positively correlated (r = 0.61, p < 2.2e-16) (Figure 2E). This suggested that the supplementation with different dosages have a very similar effect on the hamster liver. Thus, we focused on Dose II group in the following analyses as it demonstrated more distinct and significant changes associated with the supplementation of CMA. The KEGG enrichment analysis showed that 14 pathways were up-regulated after 4-week supplementation of CMA at Dose II (Figure 2C, Dataset S2). Of these, the most up-regulated pathways involved biosynthesis and metabolism of amino acids, including glycine, serine, threonine, tryptophan, alanine, aspartate glutamate, arginine and proline, as well as BCAAs (Figure 2C, Dataset S2). GO enrichment analysis of up-regulated DEGs showed that the oxoacid metabolic process, carboxylic acid metabolic process, and alpha-amino acid metabolic process are the most significantly altered biological processes (Dataset S2) in Dose II group vs HFD group.

### Reporter metabolites through the global analysis of transcriptomics data

To evaluate the detailed metabolic differences in hamsters with or without CMA treatment upon HFD feeding, we constructed a liver-specific genomic-scale metabolic model for Golden Syrian hamster based on the Golden Syrian hamster orthologs of mouse genes in mouse metabolic reaction (MMR) ^28^, and transcriptomics data by using INIT (Integrative Network Inference for Tissues) algorithm ^29^ (Figure 3A). The constructed metabolic model, namely *iHamsterHepatocyte1818*, includes 3,867 metabolic reactions, 1,818 genes, and 3,205 metabolites.

**Figure 3.**
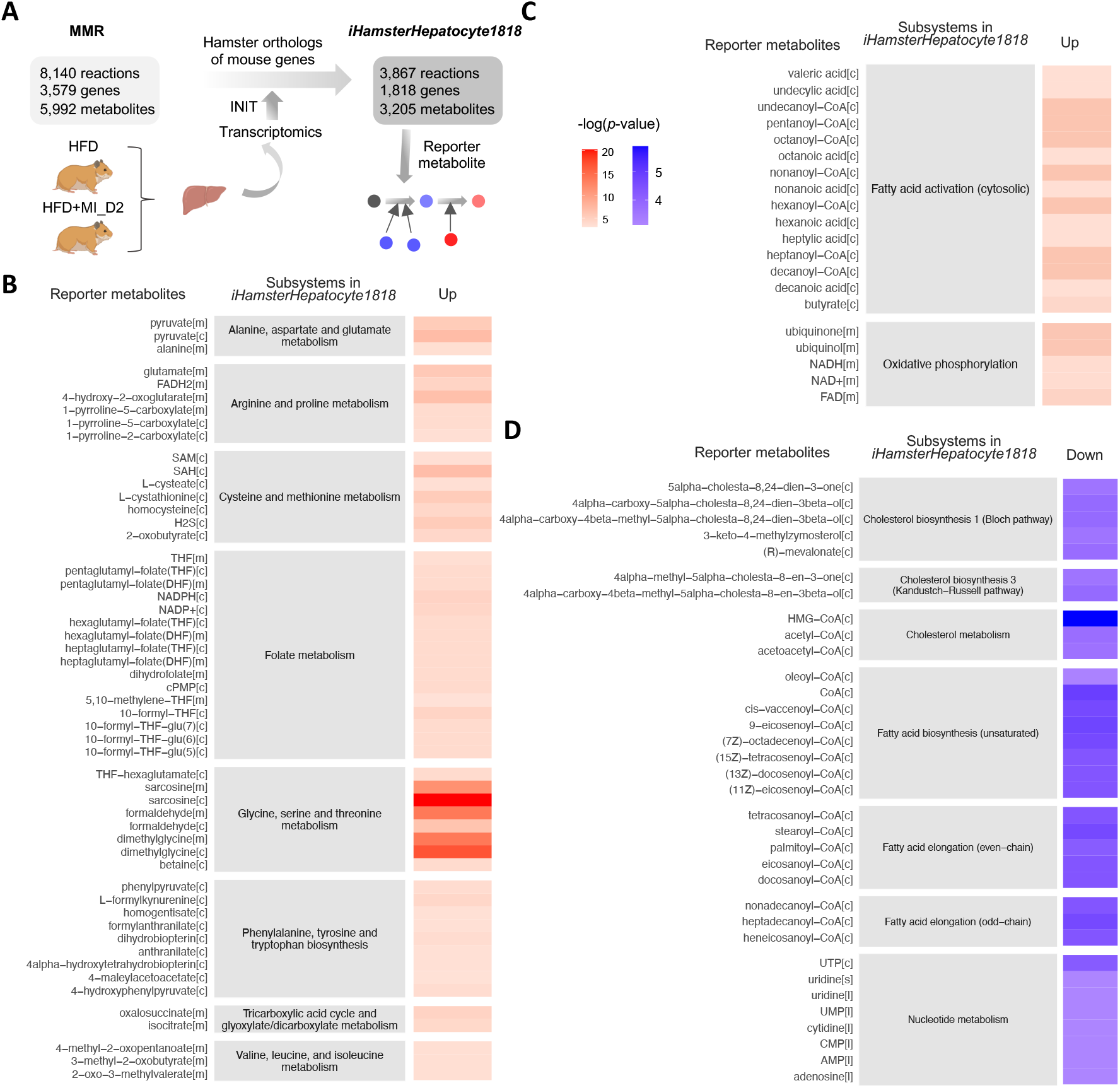
Reporter metabolites in CMA-treated group with Dose II. Metabolic differences between the hamsters treated with or without CMA at Dose II were investigated through a comparative analysis of the gene expression profiles (RNA-seq) of the liver and *iHamsterHepatocyte1818*. (a) Liver-specific genome-scale metabolic model for Golden Syrian hamster was created using the Golden Syrian hamster orthologs of mouse genes based on the MMR. (b) reporter metabolites associated with amino acid metabolism and TCA cycle; (c) fatty acid activation and oxidation phosphorylation; (d) biosynthesis and metabolism of cholesterol, biosynthesis and elongation of fatty acid, and nucleotide metabolism. p values for each reporter metabolite were calculated for up-and downregulated genes. Abbreviations: MMR, mouse metabolic reaction; INIT, Integrative Network Inference for Tissue.

Next, we performed reporter metabolite analysis to identify the key metabolic hubs in response to the supplementation of CMA with Dose II. We identified reporter metabolites using the differentially expression genes of Dose II vs HFD group and the network topology provided by *iHamsterHepatocyte1818*. Reporter metabolite analyses were used as statistical test to identify metabolites in the network for which a significant change occurred between the compared conditions. A total of 256 reporter metabolites (Reporter Features, p-value < 0.05) for hamsters treated with CMA with Dose 2 were identified (Dataset S3). Of these, 143 and 113 reporter metabolites were associated with up-regulated and down-regulated genes in CMA-treated group, respectively. The association of the reporter metabolites with up- and downregulated genes and their metabolic subsystems classified in *iHamsterHepatocyte1818* are presented (Figure 3B, C&D). As shown in the figure, CMA treatment modulated several subsystems that are known to be associated with NAFLD, including amino acid metabolism (BCAAs, cysteine, methionine, glycine, serine, threonine, arginine, proline, alanine, aspartate, and glutamate), folate metabolism, tricarboxylic acid cycle (TCA) cycle (Figure 3B), fatty acid activation and oxidative phosphorylation (Figure 3C). In addition, CMA supplementation regulated the subsystems associated with biosynthesis and metabolism of cholesterol, biosynthesis and elongation of fatty acid, as well as nucleotide metabolism (Figure 3D).

### CMA boosts hepatic metabolism to attenuate HS

Based on the integrative systems analysis, we found the CMA supplementation enhanced several NAFLD associated key metabolic pathways in the liver. We found that CMA supplementation, or more specifically, betaine supplementation, increased hepatic expression of genes involved in folate metabolism, including *SHMT2, MTHFD1, DMGDH, SARDH* and *GNMT* (Figure 4A&B). This implicated an enhanced folate cycle in the liver. Besides, we observed a significant increase in the hepatic expression of *CBS* and *CTH* involved in transsulfuration pathway, where the supplemented NAC might play an important role (Figure 4A&B). This may contribute to the GSH generation by synthesising cysteine and may play a crucial role in maintaining cellular redox homeostasis ^30^.We also observed increased hepatic expression of key genes involved in mediating GSH metabolism, including *GPX1, GPX4, GSTK1* and *PRDX5* (Figure 4A&B). Based on reporter metabolite analysis, we also identified betaine, dimethylglycine, sarcosine, 5,10−methylene−THF, 10−formyl−THF, THF, homocysteine, L-cystathionine and S-(formylmethyl) glutathione, which are associated with up-regulated genes in folate metabolism, transsulfuration pathway and glutathione metabolism (Figure 3B, Figure 4A&B). Considering that one of the significant components of the CMA is betaine which is a precursor for these metabolic pathways (Figure 4A), these results suggested that the supplementation promoted the glutathione biosynthesis through the folate cycle and transsulfuration pathway.

**Figure 4.**
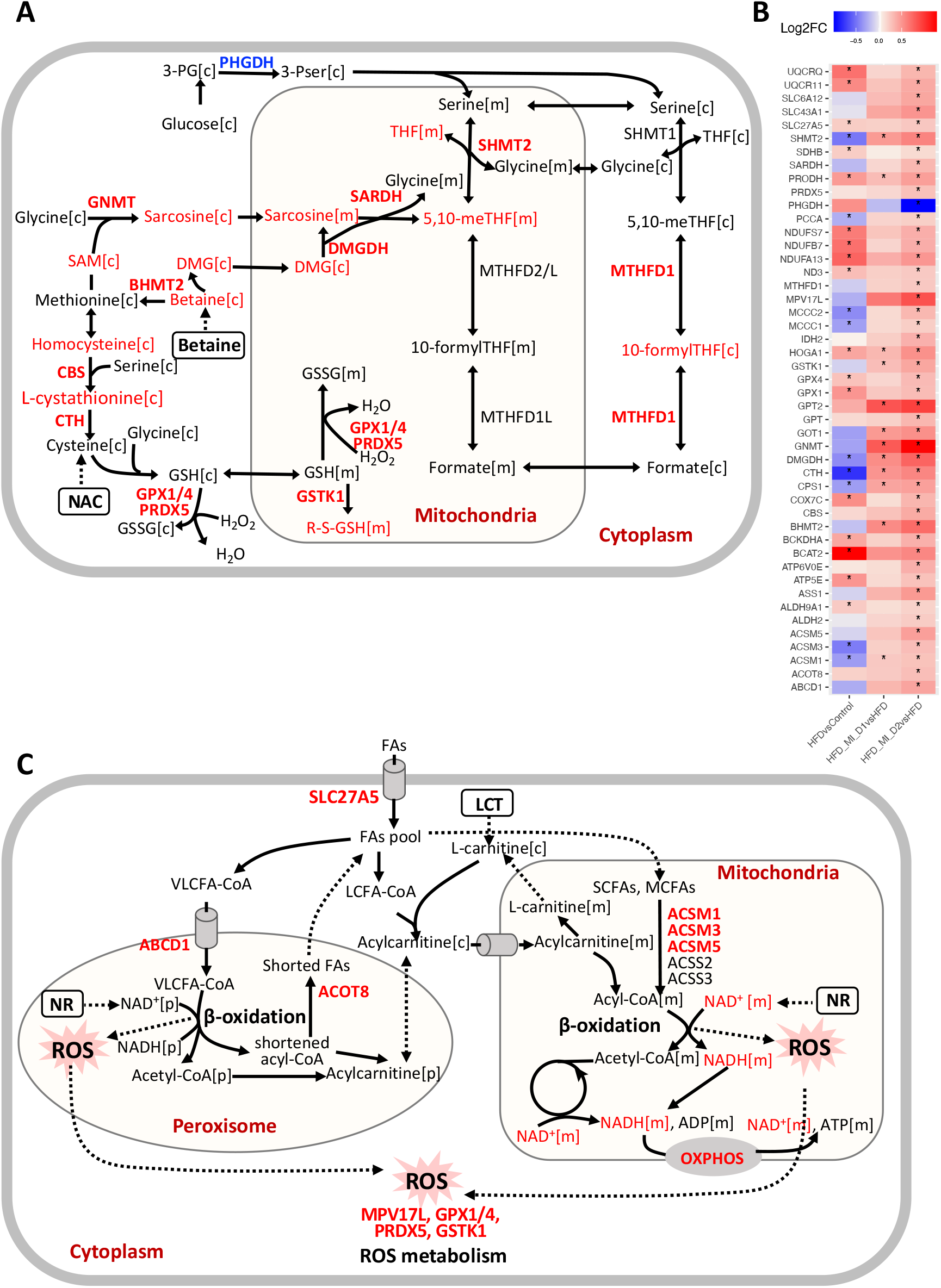
Significant increases (red) and decreases (blue) in genes and associated reporter metabolites in liver from hamsters in HFD+MI_D2 as compared to of those in HFD group. (a) Reactions involved in one-carbon metabolism and transsulfuration pathway. (b) A summary of significant diffident genes involved in one-carbon metabolism, transsulfuration pathway, fatty acid metabolism, oxidative phosphorylation (OXPHOS), BCAAs degradation, TCA cycle and urea cycle. * demonstrates the significance (adjusted p-value < 0.1). (c) Reactions involved in fatty acid metabolism in both mitochondria and peroxisome. Abbreviations: *3-PG*, 3-phosphoglycerate; *PHGDH*, phosphoglycerate dehydrogenase; *3-PSer*, 3-phosphoserine; *THF*, tetrahydrofolate; *SHMT2*, serine hydroxymethyltransferase, mitochondrial; *MTHFD1*, C1-THF synthase; *GNMT*, glycine N-methyltransferase; *DMG*, dimethylglycine; *SARDH*, sarcosine dehydrogenase; *DMGDH*, dimethylglycine dehydrogenase; *BHMT2*, S-methylmethionine--homocysteine S-methyltransferase BHMT2; *CBS*, cystathionine beta-synthase; *CTH*, cystathionine gamma-lyase; *GSH*, glutathione; *GSSG*, glutathione disulfide; *GPX1*, glutathione peroxidase 1; *GPX4*, phospholipid hydroperoxide glutathione peroxidase; *SAM*, S-Adenosyl methionine; *PRDX5*, peroxiredoxin-5, mitochondrial; *R-S-GSH*, S-(formylmethyl)glutathione; *SLC27A5*, bile acyl-CoA synthetase; *ABCD1*, ATP-binding cassette sub-family D member 1; *ACOT8*, Acyl-coenzyme A thioesterase 8; *ACSM1*, Acyl-coenzyme A synthetase ACSM1, mitochondrial; *ACSM3*, Acyl-coenzyme A synthetase ACSM3, mitochondrial; *ACSM5*, Acyl-coenzyme A synthetase ACSM5, mitochondrial; *ACSS2*, Acetyl-coenzyme A synthetase, cytoplasmic; *ACSS3*, Acyl-CoA synthetase short-chain family member 3, mitochondrial; *ASS1*, argininosuccinate synthase; *CPS1*, carbamoyl-phosphate synthase [ammonia], mitochondrial; *MPV17L*, mpv17-like protein; *GSTK1*, glutathione S-transferase kappa 1; *COX7C*, cytochrome c oxidase subunit 7C; *ATP6V0E*, v-type proton ATPase subunit e 1; *ATP5E*, ATP synthase subunit epsilon; *ALDH2*, aldehyde dehydrogenase, mitochondrial; *ALDH9A1*: 4-trimethylaminobutyraldehyde dehydrogenase; *BCAT2*, branched-chain-amino-acid aminotransferase, mitochondrial; *BCKDHA*, 2-oxoisovalerate dehydrogenase subunit alpha, mitochondrial; *GPT2*, alanine aminotransferase 2; *HOGA1*, 4-hydroxy-2-oxoglutarate aldolase, mitochondrial; *IDH2*, isocitrate dehydrogenase [NADP], mitochondrial; *MCCC1*, methylcrotonoyl-CoA carboxylase subunit alpha, mitochondrial; *MCCC2*, methylcrotonoyl-CoA carboxylase beta chain, mitochondrial; *PCCA*, propionyl-CoA carboxylase alpha chain, mitochondrial; *PRODH*, proline dehydrogenase 1, mitochondrial; *SDHB*, succinate dehydrogenase [ubiquinone] iron-sulfur subunit; *GOT1*, aspartate aminotransferase, cytoplasmic; *GPT*, alanine aminotransferase 1; *UQCRQ*, cytochrome b-c1 complex subunit 8; *UQCR11*: cytochrome b-c1 complex subunit 10; *SLC6A12*, sodium- and chloride-dependent betaine transporter; *SLC43A1*, large neutral amino acids transporter small subunit 3; *SDHB*, succinate dehydrogenase [ubiquinone] iron-sulfur subunit; *NDUFS7*, NADH dehydrogenase [ubiquinone] iron-sulfur protein 7; *NDUFB7*, NADH dehydrogenase [ubiquinone] 1 beta subcomplex subunit 7; *NDUFA13*, NADH dehydrogenase [ubiquinone] 1 alpha subcomplex subunit 13; *ND3*, NADH-ubiquinone oxidoreductase chain 3; *FAs*, fatty acids.

Peroxisome and mitochondria co-operatively perform diverse metabolic processes, including fatty acid β-oxidation and cellular ROS homeostasis ^31^. Short- and medium-chain fatty acids (SCFAs, MCFAs) are oxidized in the mitochondria whereas long-chain fatty acids (LCFAs) are oxidized in both mitochondria and peroxisome. Very-long-chain fatty acids (VLFAs) are preferentially oxidized in the peroxisomes ^32^. We found that the hepatic expression of genes associated with fatty acid oxidation in peroxisome and mitochondria (*SLC27A5, ABCD1, ACOT8, ACSM1, ACSM3 and ACSM5*) were elevated in the liver of CMA-treated hamsters, which could be related to the supplementation of LCT and NR (Figure 4B&C). Besides, we observed the genes involved in ROS metabolism including *MPV17L, GSTK1, PRDX5, GPX1*, and *GPX4* (Figure 4B&C), showed a significant increase which could be a result of the supplemented NR and elevated glutathione metabolism. *SLC27A5*, which facilitates the uptake of LCFAs, has been considered as long-chain and very-long-chain acyl-CoA (VLCFA-CoA) synthetases ^33, 34^. Interestingly, the hepatic expression of genes involved in *de novo* synthesis of NAD^+^, an important coenzyme for redox reactions, including *TDO2, AFMID* and *KYNU* were significantly increased after CMA supplementation. NAD^+^ and NADH were also identified as reporter metabolites associated with upregulated genes in fatty acid oxidation, oxidative phosphorylation and TCA cycle (Figure 4B&C). Notably, in agreement with the aforementioned findings, we found that NAD^+^ level was significantly increased in the liver with CMA Dose I and II supplementation vs HFD (Figure 1J).

Additionally, we found that CMA supplementation increased the hepatic expression of genes involved in BCAAs catabolism, including *BCAT2, BCKDHA, MCCC1, MCCC2, ALDH2, ALDH9A1* and PCCA (Figure 4B, Figure 5A). We also observed significantly enhanced hepatic expression of *IDH2* and *SDHB* in TCA cycle. Increased TCA cycle and BCAAs catabolism could indicate that the fatty acid oxidation and cellular respiration are enhanced as suggested by a previous study ^35^. Moreover, we observed that the hepatic expression of genes involved in arginine biosynthesis (*GPT, GOT1, GPT2, CPS1* and *ASS1*) and genes involved in arginine and proline metabolism were significantly increased with the CMA supplementation. It has been reported that the downregulation of *CPS1*, the flux-generating urea cycle feeder enzyme, correlated with the loss of functional capacity for ureagenesis in patients with NASH ^36, 37^.

**Figure 5.**
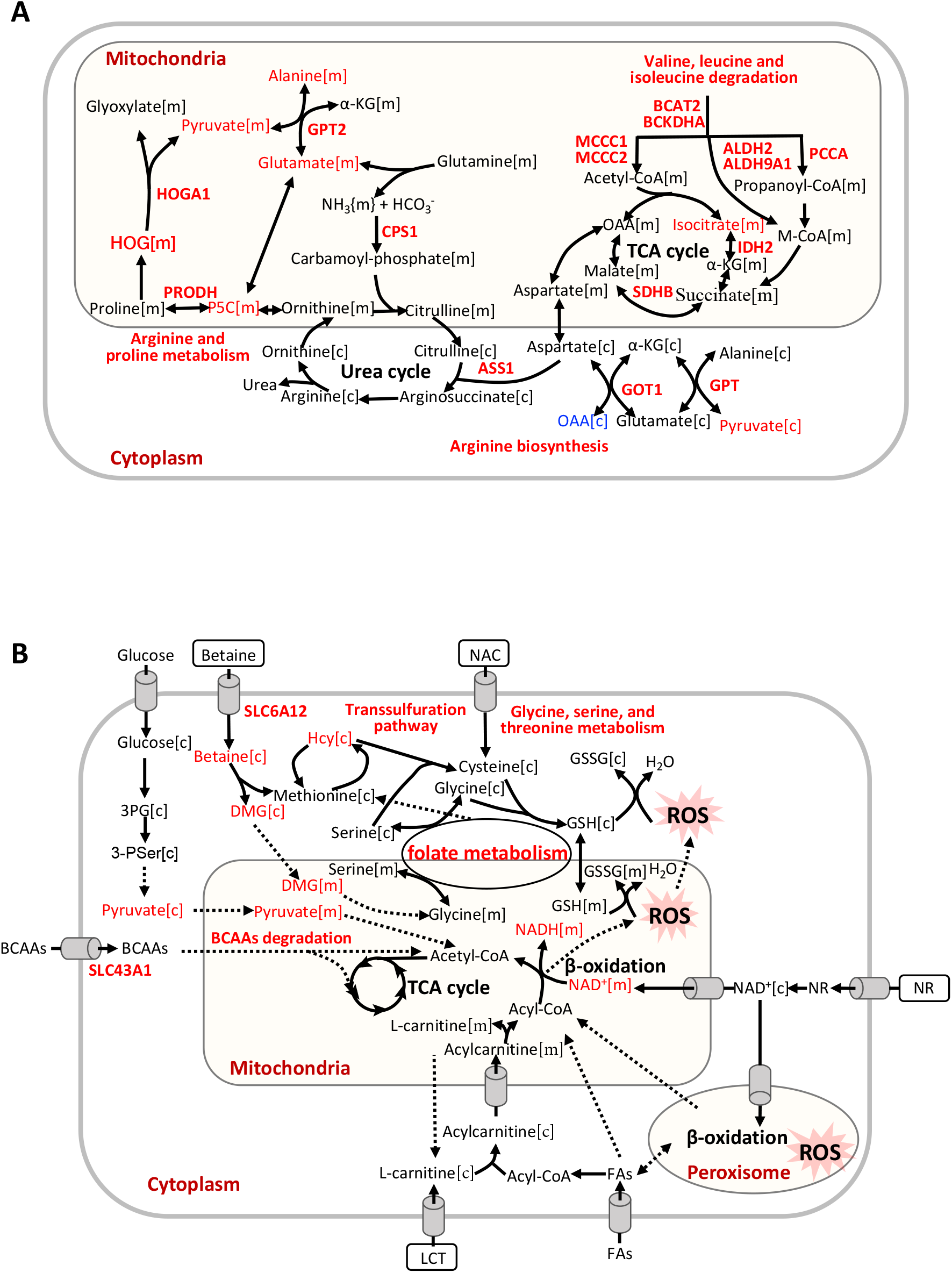
(a) Reactions involved in BCAAs degradation, tricarboxylic acid cycle, and urea cycle. (b) Global effect of CMA supplementation. Abbreviations: *OAA*, Oxalacetic acid; *α-KG*, alpha-ketoglutarate; *HOG*, 4−hydroxy−2−oxoglutarate; *M-CoA*, methylmalonyl-CoA; *P5C*, 1−pyrroline−5−carboxylate; *Hcy*, homocysteine, *NR*, nicotinamide riboside; *NAC*, N-acetyl-l-cysteine; *FAs*, fatty acids; *LCT*, L-carnitine tartrate.

## Disscussion

In this study, we evaluated the global changes induced by the CMA treatment (Figure 5B). We demonstrated that HFD-fed hamsters that received the CMA at both doses displayed an amelioration of hepatic steatosis compared to their HFD-fed control counterparts. This finding was accompanied by a significant decrease in final body weight and body weight gain in the hamsters supplemented with the Dose II, which can be attributed to the lower caloric intake observed in these animals. Interestingly, our transcriptomics analysis revealed that after CMA treatment there was a significant increase in the hepatic expression of genes in one-carbon metabolism and transsulfuration pathway, which are closely associated to the *de novo* GSH synthesis, which, in turn, is altered in patients with HS ^15^. Moreover, we reported that CMA supplementation increased the expression of genes involved in BCAAs catabolism and urea cycle in the liver. Furthermore, we observed elevated NAD^+^ hepatic levels, which implicated the enhancement of multiple metabolic pathways including fatty acids oxidation, the lack of which has a critical role in the development of NAFLD ^15, 38^, and strongly suggest CMA supplementation as a very promising strategy to ameliorate fatty liver.

Our findings agreed well with and also extended the predicted results from our recent work ^20^. Our previous studies revealed that altered GSH and NAD^+^ metabolism is a prevailing feature of NAFLD ^15^. Amino acids, in particular serine, glycine, methionine, and several metabolites involved in one-carbon metabolism play crucial roles in NAFLD progression ^14^. Mice with HFD-induced NAFLD also showed dysregulated one-carbon metabolism in liver ^39, 40^. Dysregulated BCAAs metabolism is significantly associated with the progression of NAFLD ^41^. The increased plasma level of BCAAs have been reported in NAFLD metabolomics studies ^42^. Based on our previous study, we predicted that catabolism of BCAAs was significantly increased after CMA supplementation in based on integration of GEMs and metabolomics data ^20^.

Serine hydroxymethyltransferase (*SHMT)* catalyses the interconversion of glycine and serine, which are precursors for the generation of GSH, an important antioxidant for maintaining the redox balance in fatty acid β-oxidation ^15, 43^. Glycine *N*-methyltransferase (*GNMT*), the main gene involved in liver S-adenosylmethionine (SAM) catabolism, is down-regulated in the liver of HFD hamsters ^44^, as well as patients with cirrhosis and HCC ^45^. In other studies, deletion of *GNMT* in mice led to excess increase in hepatic SAM, which is associated with fatty acid metabolism, oxidative stress and HS ^46, 47^. The down-regulation of *DMGDH*, which is a mitochondrial dimethylglycine dehydrogenase, was associated with the insulin resistance in a previous study ^48^. Hepatic *CBS/CTH* system was even positioned as a potential therapeutic target in NAFLD due to the deficiencies of *CBS* and *CTH* in rodents’ study ^49, 50^. Taking into account the aforementioned findings ^51^, in the present study, the increased expression of key genes involved in folate metabolism (*SHMT2, MTHFD1, DMGDH, SARDH*, and *GNMT*) and in the transsulfuration pathway (*CBS* and *CTH*) found in the livers of CMA-supplemented hamsters strongly suggest that CMA ameliorated NAFLD promoting glutathione biosynthesis through the activation of these two metabolic pathways.

*IDH2* is considered a unique enzyme in the regulation of mitochondrial ROS ^52^. *IDH2* knockout accelerates oxidative stress, lipid accumulation and HS in HFD-challenged mice, which were restored by promoting *IDH2* expression ^53^. In the present study, we found that *IDH2* was upregulated in the livers of hamsters after CMA supplementation vs HFD. Relevantly, the ROS metabolism-related genes *MPV17L* and *GSTK1* displayed the same pattern of expression than those observed for *IDH2. MPV17L*, a transmembrane protein, has been implicated in the regulation of peroxisomal ROS metabolism ^54^. *MPV17L* also prevents mitochondrial dysfunction and apoptosis through its antioxidant and antiapoptotic properties *in vivo* and *in vitro* ^55^. *GSTK1* is a highly conserved enzyme potentially involved in redox reaction and has a crucial role in protection against oxidative stress ^56^. In addition, the expression of *GPX1* and *GPX4* genes, which codify for the glutathione peroxidase (GPx) family of enzymes, were also up-regulated in the hamsters that received the CMA treatment. *GPx* catalyses the reduction of H_2_O_2_ and lipid peroxides through the conversion of GSH to GSSG and thereby plays a crucial role in the protection of cells against oxidative damage. Patients with NASH had significantly lower levels of *GPx* activity ^57^. Overall, our results strongly suggest that CMA would protect against NAFLD-associated oxidative stress, at least in part, through the up-regulation of the expression of these key genes involved in ROS metabolism.

The increased hepatic expression of genes associated with fatty acid oxidation in peroxisome and mitochondria (*SLC27A5, ABCD1, ACOT8, ACSM1, ACSM3* and *ACSM5*) found in response to the supplementation with CMA suggest that this multi-ingredient would tackle NAFLD by boosting this catabolic pathway in both organelles. Different previous findings contribute to reinforce our hypothesis. Thus, a recent study showed that knockout of *SLC27A5* lead to the disrupted lipid metabolism and redox balance in HCC cells both *in vitro* and *in vivo* ^58^. In patients with severe steatohepatitis and cirrhotic liver, the hepatic expression level of *SLC27A5* were significantly decreased and inversely correlated with histological progression ^59^. *ABCD1* mediates the uptake of the VLCFAs across the peroxisome membrane and its loss of function results in defective β-oxidation of VLCFA and increased cellular oxidative stress ^60, 61^.

In conclusion, the supplementation for 4 weeks with a cocktail of metabolic activators (CMA) including L-carnitine, N-acetyl-l-cysteine, nicotinamide riboside and betaine at 2 intended human clinical doses ameliorated HS in HFD-fed Golden Syrian hamsters. This health effect was accompanied by body weight loss and decreased caloric intake in the animals that received Dose II. To shed light on the metabolisms by which CMA exerted its effects, we performed liver transcriptomics analysis and analysed the data using GEMs. We observed that CMA supplementation significantly attenuated the HFD-induced HS by modulating the GSH and NAD^+^ metabolisms, as well as promoting fatty acid oxidation, BCAAs catabolism and urea cycle and eventually activating mitochondria. Our findings provide extra evidence about the beneficial effects of a treatment based on these metabolic activators against NAFLD.

## Methods and materials

### Animal model

The Animal Ethics Committee of the Technological Unit of Nutrition and Health of Eurecat (Reus, Spain) and the *Generalitat de Catalunya* approved all procedures (DAAM 10026). The experimental protocol complied with the ARRIVE guidelines, followed the ‘Principles of laboratory animal care’ and was carried out in accordance to the EU Directive 2010/63/EU for animal experiments. All animals were housed individually at 22 °C under a light/dark cycle of 12 h (lights on at 09:00 am) and were given free access to food and water.

Thirty-nine 10-week-old male Golden Syrian hamsters (Janvier Labs, Saint Berthevin, France) weighting 110-120 g were used. After an adaptation period of 1 week, hamsters were randomly assigned into two experimental groups fed with a NFD (n = 9, 11% calories from fat; Envigo, Barcelona, Spain) or a HFD (n = 30, 23% calories from fat and 1g/kg cholesterol; Envigo, Barcelona, Spain) (Dataset S4) for 8 weeks. In a previous study, using very similar diets we demonstrated that this period was useful to induce fatty liver and hypercholesterolemia in hamsters ^25^. Afterwards, HFD group was further randomly distributed into three subgroups: placebo, CMA at Dose I (200mg/kg LC, 200 mg/kg NAC, 200 mg/kg NR and 400 mg/kg betaine; HFD+MI_D1 group) and multi-ingredient at Dose II (400mg/kg LC, 400 mg/kg NAC, 400 mg/kg NR and 800 mg/kg betaine; HFD+MI_D2 group) for the last 4 weeks. LCT contained 68.2% of LC and, therefore, to reach the desired LC doses hamsters were supplemented with 293 mg/kg and 586 mg/kg of LCT for Dose I and Dose II, respectively. LCT, NAC and NR were daily diluted with low-fat condensed milk diluted 1:3 with water (vehicle) and orally given to the hamsters. Four days before the beginning of the treatments, the animals were trained to lick diluted low-fat condensed milk diluted 1:3 with water (0.2 mL) to ensure voluntary consumption. It was confirmed that each hamster fully ingested the daily dose of the corresponding treatment, which was given daily beween 8:30-10:00 am. Betaine was included in opaque bottles containg the drinking water (4g/L and 8g/L fos Dose I and Dose II, respectively) and renewed three times per week. Considering an average hamster’s weight of 125g, the doses of LCT, NAC and NR used were equivalent to the daily consumption of 1,918 mg and 3,836 mg of these metabolic activators for a 60-kg human ^62^, for the Dose I and Dose II, respectively. For betaine, the extrapolated daily intake using the same formula were 3,836 mg and 7,672 mg. These dosages are considered acceptable and safe in a context of a multi-ingredient supplementation to tackle NAFLD ^19^.

Body weight and food intake were recorded once per week, and food was renewed daily. At 12 weeks, all experimental animals were sacrificed under anaesthesia (pentobarbital sodium, 60 mg/kg body weight) after 6 h of diurnal fasting. Blood was collected by cardiac puncture, and serum was obtained by centrifugation and stored at -20 °C until analysis. The liver, soleus and gastrocnemius muscles, and white adipose tissue (WAT) depots (retroperitoneal (RWAT) and mesenteric (MWAT) depots) were rapidly removed, weighed, frozen in liquid nitrogen and stored at -70 °C until analysis.

### Histological evaluation

Morphometric analyses of tissues and steatosis of liver histology were described as before ^25^. In general, the entire histological section (approximate area 2 cm^2^) of the liver were analysed according to Brunt’s golden standard score, estimating the percentage of the area covered by fat droplets and using a scored from 0 and 3 (0, null, 1, when steatosis was detected in up to 30% of the area; 2, when steatosis was observed in between 30 and 60% of the area; 3, when steatosis was observed in more than 66% of the area).

### Body Composition Analyses

Body composition were analysed by Nuclear magnetic resonance (NMR). Lean and fat mass analyses were performed at the end of weeks 8 and 12 using an EchoMRI-700^®^ device (Echo Medical Systems, L.L.C., Houston, TX, United States). The measurements were performed in duplicate. Data are expressed in relative values as a percentage of body weight (%). Lean/fat mass ratio was also calculated.

### Serum Analysis

Enzymatic colorimetric assays were used for the analysis of glucose, total cholesterol and triglycerides (QCA, Barcelona, Spain), HDL-cholesterol and LDL/VLDL-cholesterol (Bioassay systems, California, USA). Serum insulin levels were analysed using a hamster insulin ELISA kit (MyBiosource, Bizkaia, Spain).

### ^1^H NMR analysis for NAD^+^ determination

For ^1^H NMR analysis, the metabolite aqueous extracts obtained in the liver were reconstituted in 700 ul of a solution containing trisilylpropionic acid (TSP) (0.74 mM) dissolved in D_2_O phosphate buffer (0.05 M). Samples were vortexed, homogenised for 5 min and centrifuged (15 min at 14000 x g). Finally, the redissolved extractions were transferred into 5 mm NMR glass tubes. ^1^H NMR measurements were performed following the procedure described by *Vinaixa* et al. (2010) ^63^.

### Transcriptomics and Gene Set Enrichment analysis

Total RNA was extracted from the liver samples using TriPure reagent (Roche Diagnostic, Sant Cugat del Valles, Barcelona, Spain) according to the manufacturer’s instructions, and used in the subsequent mRNA sample preparation for sequencing using NovaSeq 6000 S1 platform with the standard Illumina RNA-seq protocol. The RNA-seq was performed at National Genomics Infrastructure (NGI, Sweden) (project number P15763). Gene abundance (in both raw counts and transcripts per million) was quantified using the kallisto pipeline ^64^ based on Golden Syrian hamster (*Mesocricetus auratus*) genome (version MesAur1.0.101). In order to validate the intra-group homogeneity, we first performed a principal component analysis (PCA). We filtered out the S105 sample in control group based on the results of PCA analysis. Next, we used DESeq2 package in *R* ^65^ to identify differentially expressed genes (DEGs) (adjusted p value < 0.1). Gene Ontology and KEGG pathway analysis were performed separately on upregulated and downregulated genes using DAVID Bioinformatics Functional Annotation Tool ^66^. Only ‘biological process’ categories enriched with a BH-corrected *P* ≤ 0.05 were considered for GO category analysis.

### Metabolic model reconstruction and Reporter metabolite analysis

To provide a resource for automated and semi-automated reconstruction of liver-specific GEMs for Golden Syrian hamster, we constructed *iHamsterHepatocyte1818* based on the MMR databases, transcriptomics data, and INIT (Integrative Network Inference for Tissues) algorithm, which has been used to reconstruct the functional GEMs based on bulk transcriptomics data as well as 56 user-defined metabolic tasks, which are known to occur in cells/tissues ^67^ and should be performed by the resulting metabolic model. A reporter metabolite analysis was performed using RAVEN toolbox implemented in *MATLAB* 2020a based on DEGs obtained as described above, together with *iHamsterHepatocyte1818*.

## Statistical analysis

The continuous variables of biological assay were showed as mean ± SD. Grubbs’ test was used to detect outliers, which were discarded for subsequent analyses. Differences between two groups were analysed by *t*-test from the *SciPy* package. Differences in steatosis scores were detected using the Chi-squared test. Missing values were dropped before the analysis. Categorical variables were analysed using Chi-square test. Principal component analysis (PCA) was performed on transcriptomics data sets to explore the quality of data and detect possible outliers. All results were considered statistically significant at *p* < 0.05. Statistical analyses were performed using Python 3.7 and *R* (v4.0.3) was used for data visualisation.

## Conflict of interest statement

AM, JB and MU are the founder and shareholders of ScandiBio Therapeutics and they filed a patent application on the use of CMA to treat NAFLD patients. The other authors declare no conflict of interest.

## Acknowledgments

We gratefully acknowledge the help of Yaiza Tobajas, Anna Antolín, Iris Triguero, Cristina Egea and Gertruda Chomiciute, who are laboratory technicians at the Technological Unit of Nutrition and Health of Eurecat, for their technical support. The author would like to thank NGI Sweden for generation of transcriptomics data. AM and HY acknowledge support from the PoLiMeR Innovative Training Network (Marie Skłodowska-Curie Grant Agreement No. 812616) which has received funding from the European Union’s Horizon 2020 research and innovation programme. The computations were performed on resources provided by SNIC through Uppsala Multidisciplinary Center for Advanced Computational Science (UPPMAX) under Project SNIC 2020/16-69.

## Funding

This work was financially supported by the Agency for Business Competitiveness of the Government of Catalonia (ACCIÓ) [TECCT11-1-0012]. This work was financially supported by the Centre for the Development of Industrial Technology (CDTI) of the Spanish Ministry of Science and Innovation under grant agreement: TECNOMIFOOD project (CER-20191010). This work was financially supported by ScandiBio Therapeutics and Knut and Alice Wallenberg Foundation.

## Author contributions

A.M., A.C., M.U., and J.B. designed the study; J.M.P., N.B., J.M.B., and L.A. did the experiments; H.Y., M.Y., and C.Z. analyzed the transcriptomics data; H.Y. and C.Z. interpreted the data; H.Y. wrote the manuscript; and all authors helped revise the manuscript.

## Supplementary datasets

Dataset S1: List of genes identified in liver transcriptomics and DEGs comparing HFD to control, HFD+MI_D1 to HFD, HFD+MI_D2 to HFD, and HFD+MI_D1 to HFD+MI_D2, respectively.

Dataset S2: KEGG pathway and GO enrichment analysis for DEGs in HFD versus control and HFD+MI_D2 versus HFD.

Dataset S3: Reporter metabolites are presented using the differential expression pf genes in the liver tissue of the CMA-treated hamsters compared with that of the liver tissue of the HFD hamsters.

Dataset S4: Composition of the diets used in the study.

